# Emergence of ST3390: A non-pigmented HA-MRSA clone with enhanced virulence

**DOI:** 10.1101/2025.03.18.643911

**Authors:** Emily A. Felton, Mary-Elizabeth Jobson, Ariana M. Virgillio, Joshua Alvior, Eleonora Cella, Amorce Lima, Deanna Becker, Suzanne Silbert, Taj Azarian, Kami Kim, Lindsey N. Shaw

## Abstract

**Background:** One of the most successful and widely-distributed hospital-associated lineages of MRSA is CC5. These strains are known from widespread antibiotic resistance, but less severe disease than CA-MRSA counterparts. Recently, CC5 descendant lineages have appeared globally with hypervirulent properties.

**Methods:** Herein we use genomic analyses to study the epidemiology of a rare CC5 MRSA sequence type, ST3390, circulating within Tampa General Hospital (TGH). We employ genetic tools alongside *in vitro* and *in vivo* models of virulence to study the pathogenic capabilities of strains.

**Results:** To date, there have only been 50 recorded instances of infection caused by ST3390 globally, with 36 of those occurring at TGH. Genomic analysis of strains identified numerous *spa*-types, with a t010 cluster found only at TGH. Exploration of AMR genes detected the presence of unique hybrid SCCmec types, with ∼90% of TGH strains possessing components of SCCmecIa, SCCmecIIa and/or SCCmecVIII. Phenotypically, all ST3390 strains lack the staphyloxanthin pigment, which is mediated by a conserved 6 aa in frame deletion within the staphyloxanthin biosynthesis protein CrtN. ST3390 strains display high levels of cytotoxicity towards human neutrophils compared to other CC5 lineages, with several isolates displaying hypervirulence in animal models of infection.

**Conclusions:** This is the first study to characterize the pathogenicity and genomic architecture of the rare MRSA lineage ST3390. Our work provides a deeper understanding of the clonal expansion of CC5, and the wider diversification of *S. aureus* isolates within patient populations.

## Introduction

Methicillin-resistant *Staphylococcus aureus* (MRSA) is an important cause of antibiotic-resistant infections that range in severity from localized skin and soft tissue infections to invasive bacteremia. Over the last 30 years MRSA infections have undergone an epidemiological shift, transitioning from a largely hospital associated problem (HA-MRSA) to also infecting young healthy individuals within the community (CA-MRSA) [1, 2]. To aid in our understanding and tracking of MRSA infections, tools have been developed for typing strains. These include variations in the repeat region of the staphylococcal protein A (*spa*) [3], and in the Staphylococcal cassette chromosome mec (SCCmec) [4]. Finally, perhaps the most important typing system is through 7 housekeeping genes that are used for multi locus sequence typing (MLST). Strains that share 6 MLST sequences are grouped into the broader classification of clonal complex (CC); with each CC comprising multiple closely related sequence types (ST) [5].

One of the most important HA-MRSA lineages is CC5 and its subtype ST5 strains. CC5s possess SCCmec type elements that are larger (I, II and III) and confer resistance to multiple classes of antibiotics [6]. Conversely, the most common CA-MRSA lineage is CC8-ST8, strains of which commonly have smaller SCCmec types (IV) and mobile elements that carry toxins such as the Panton-Valentine Leukocidin [7]. The lines separating HA-MRSA and CA-MRSA strains, however, have blurred within the last decade. Recent studies have illustrated the evolution of CA-MRSA becoming a leading cause of hospital-associated bloodstream infections [8, 9], whilst there have been increasing accounts of HA-MRSA strains circulating in the community or eliciting serious disseminated infections [10, 11].

Along these lines, over the last 20 years we have seen the emergence of hypervirulent HA-MRSA CC5 variant lineages. One such descendant, ST764 from Japan, has presented not only CA-MRSA characteristics but also acquisition of non-staphylococcal mobile elements [12]. Another variant, ST105, replaced ST5 as the leading cause of blood stream infections in Brazil [13, 14]. In the United States the New York/Japan clone, ST5/SCCmecII (USA100) has been the most dominant HA-MRSA lineage. Herein we present characterization of a CC5 descendant clone in North America, ST3390, that demonstrates hypervirulent characteristics as well as evidence of horizontal gene transfer with CA-MRSA strains. Collectively, our work provides a deeper understanding of the emerging clonal expansion of CC5, and the wider diversification of *S. aureus* isolates within patient populations.

## Methods

### Bacterial Strains and Growth Conditions

MRSA TGH-ST3390 strains were isolated from patients and collected by Tampa General Hospital between 2017 and 2024 [15]. All bacterial strains and plasmids used are listed in **Supplemental Table 1**. Overnight cultures were routinely grown at 37°C with shaking at 250 rpm in 5mL of tryptic soy broth (TSB). Where required, media was supplemented with the following antibiotics: for *E. coli*: 100μg/mL ampicillin; for *S. aureus*: 10μg/mL chloramphenicol, 5μg/mL erythromycin and 25μg/mL lincomycin.

### Genomic Analyses

All sequencing and assembly methods are contained in the companion paper to this work [15]. Genomes were sequence-typed with pyMLST [16] , annotated with prokka, and AMR genes identified using AMRFinderPlus v3.10.45 . The DNA sequence and translated proteins for genes of interest were confirmed and aligned using BLASTncbi-BLAST v2.13.0+. The staphtyper workflow as part of Bactopia v3.0.1 [17] was used to characterize SCCmec and *spa*-type. Genomes were analyzed for SNPs against the well-characterized CC5-ST5 strain N315 using Snippy (v4.6.0). Snippy was used to generate an initial core genome single-nucleotide polymorphism alignment. Following this, Gubbins was used to create a recombination masked alignment. From this alignment RAxML was used to infer a maximum-likelihood mid-point rooted tree. The subsequent tree was visualized and annotated using iTOL webserver. Genome regions were aligned to reference sequences and visualized with clinker. Amino acid sequences were aligned using muscle and viewed in snapgene. Prophage regions were annotated with Phastest webserver [18]. TGH-ST3390 blast maps were generated using the CGview comparison tool[19].

### *crt* Operon complementation

The *crtOMNPQ* operon was PCR amplified from TGH993 using primers OL7816 (cagtcaGTCGACgtgactacaactgcagcg) and OL7817 (cagtcaGGATCCgactactcccttatacgcccc). This fragment was cloned into pMK4 via BamHI and SalI restriction sites, creating pEAF1. The construct was transformed into *E. coli* DH5α, with clones confirmed using OL7816/OL7817, followed by Sanger sequencing (Azenta). The plasmid was then transformed into *S. aureus* RN4220 by electroporation, followed by phage transduction in a JE2 *crtN* transposon mutant [20] using φ11. Strains were grown on TSA and observed for pigmentation.

### Blood viability Assay

These were performed as described previously [21]. Bacterial cultures were grown in biological triplicate overnight before being diluted 1:100 and grown for 3h. These cultures were then washed twice with PBS before being used to inoculate 1mL of human, gender-pooled whole blood (BioIVT) at a final OD_600_ of 0.05. Samples were incubated for 6h with rotation at 37°C. Aliquots were withdrawn, serially diluted and plated to determine viability.

### Cytolytic Assays

Hemolysis, proteolysis and cytolysis assays were all performed as described [22, 23]. For proteolysis and hemolysis, these were monitored using casein- and blood-agar, respectively. Cultures were grown overnight on TSA and individual colonies were patched onto either casein agar (TSB containing 5% nonfat dry milk) or 5% sheep blood agar (Thermo Scientific). After 18h plates were imaged and quantified using ImageJ. For cytolysis, the immortalized HL-60 cell line was grown to confluence in RPMI supplemented with 10% FBS and 100U Pen/Strep at 37°C and 5% CO_2_. Cells were differentiated into neutrophil-like cells by the addition of 1.25% DMSO for 4 days; which was confirmed via cell morphology. Following this, cells were resuspended in RPMI with 10% FBS and 1x10^5^ cells were seeded in 100µl into a 96-well plate. Bacterial cultures were grown as above and standardized to OD_600_ 0.05. Seeded neutrophils were intoxicated in technical duplicate with 5µl of bacterial supernatant (5% total volume) followed by incubation at 37°C, 5% CO_2_ for 1h. Cell viability was measured using the CellTiter96 Aqueous One Solution Cell proliferation reagent (Promega). Following 1h, 20µl of the CellTiter reagent was added to wells, and the plate returned to the incubator for 2h. Color development was assessed via OD_490_ using a Biotek Cytation 5 plate reader. Data is reported as percent death of supernatant-treated cells compared to neutrophils treated with 5µl TSB.

### Murine Model of Sepsis

All animal studies in this work were performed in accordance with and approved by the Institutional Animal Care and Use Committee of the University of South Florida. To prepare inocula, bacteria was washed 2x with PBS before being adjusted to 1x10^9^ CFU/mL in PBS. Female, six-week-old CD-1 mice (Charles River Laboratories) were injected in the tail vein with 100µl of bacteria, resulting in a final dose of 1x10^8^. Infections were allowed to progress for 7 days, or until mice reach a premoribund state, at which point they were euthanized. Any animal remaining at the end of the infection period were also euthanized. For those animals surviving the 7 days, the liver and kidneys were harvested for bacterial load determination. Experiments were performed with 9 mice per strain and a Log Rank or Mann-Whitney test was used to measure statistical significance.

## Results

### Geotemporal Distribution of ST3390 Clones

In a companion study we identify 36 isolates from the rare *S. aureus* Sequence Type ST3390 at a large urban hospital in Tampa, Florida (Tampa General Hospital, TGH) [15]. To understand the relevance of this, we reviewed existing literature and available genomic databases to understand the wider distribution of this clone (**Table 1**). In so doing we found an additional 47 isolates of ST3390 worldwide, with 45 available genomes. The earliest documented infection from this lineage is a single strain from Canada in 2010[24]. From the years 2013-2016 four individual isolates caused infections across different cities within the Northeastern USA[25, 26]. Between 2015 and 2016 the University of Illinois at Chicago collected 30 ST3390 isolates from environmental surfaces and 4 isolates from 2 patients (1 was a nasal sample)[27]. Also in 2016, two nasal surveillance isolates were reported in Baltimore, MD. In 2017 multiple isolates were collected from infections in our hospital in Tampa, as well as the solitary patient isolate from outside North America in the Netherlands[28]. In 2018, one isolate was found from an infection in Jacksonville, FL. During 2019, we found 13 isolates from patients in our hospital, and 2 more were recovered from infections in Gainesville, FL. After 2021, Tampa is the only city to report ST3390 strains from infected patients. In sum, to date there have been 50 recorded instances of infection caused by ST3390 globally, with 36 of those occurring at TGH. Interestingly, when one reviews genomic relatedness of strains (**Figure 1**), the oldest isolate from Canada appears to root the tree and is similar to early strains isolated in New York. Beyond this, our TGH strains cluster relatively well with other isolates from Florida locations, and the broader population, with the exception of the Chicago strains, which cluster separately and are clearly part of an isolated outbreak.

**Figure 1.**
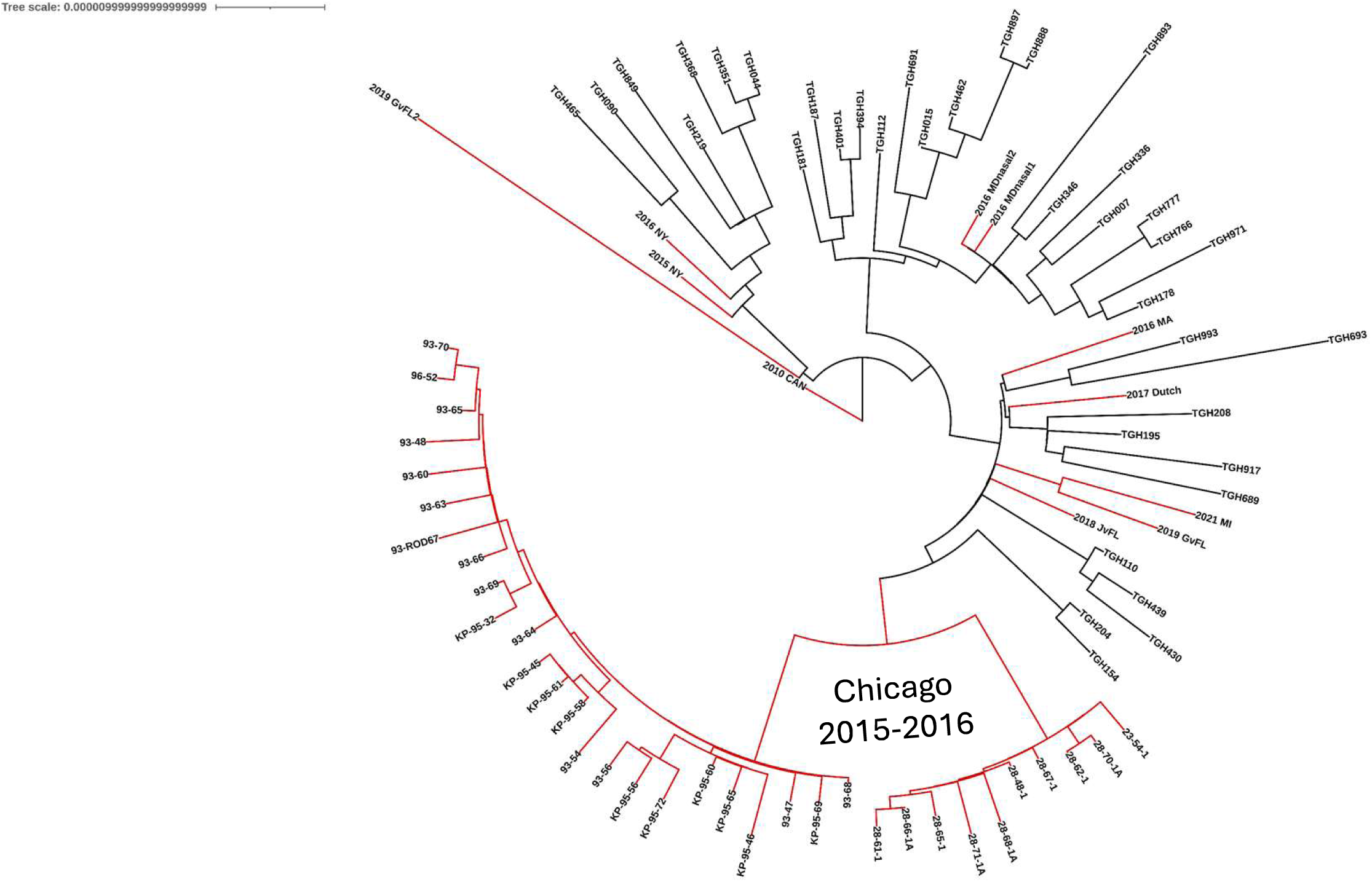
The Genomic Relatedness of ST3390 strains. Shown is a recombination masked core SNP alignment maximum-likelihood tree of the 36 TGH-ST3390 strains and 45 non-TGH ST3390 strains whose genomes are available. SNP alignment was generated with snippy v.4.6.0. Non-TGH branches are denoted in red and are named by city and year of isolation. TGH strains are shown in black.

**Table 1.** Geotemporal Distribution of ST3390 Isolates Globally. SCCmec type information for strains from this study is available in supplemental table 1. #Genomic data was either not available or was in an unusable form.

| Year | Geographic location | Number of Isolates | Spa Type | Accession | SCCmec | References |
| --- | --- | --- | --- | --- | --- | --- |
| **2010** | CNISP Ontario, Canada | 1 | t579 | SAMN13571250 | SCCmecIIa | [22] |
| **2013** | University of Chicago, Chicago, Illinois | 1 |  | # |  | pubMLST |
| **2015** | Mount Sinai Hospital, New York, NY | 1 | t002 | SAMN09484063 | SCCmecIIa |  |
| **2015-2016** | RUMC, Chicago, Illinois | 34 | t002, t062, t045 t535, t2597, t539, t111, t2379, t586, t458, t2235, t2302 | PRJNA595570 | orfX-JI -ACME-SCCmecIIa (n=10) | [25] |
| **2016** | Massachusetts | 1 | t002 | SAMN12325109 | orfX-ACME-SCCmecIIa | [23, 24] |
|  | Weill Cornell New York Presbyterian, New York, NY | 1 | t002 | SAMN12776173 | SCCmecIIa |  |
|  | University of Maryland, Baltimore, MD | 2 | t002, t045 | PRJNA565025 | orfX-JI -ACME-SCCmecIIa (n=2) |  |
| **2017** | Tampa General Hospital, Tampa, FL | 8 | t002(n=3), t062(n=2), t010(n=3) | This Study |  |  |
|  | RIVM, Netherlands | 1 | t2065 | SAMN30843088 | orfX-JI -ACME-SCCmecIIa | [26] |
| **2018** | Tampa General Hospital, Tampa, FL | 3 | t002, t062, t010 | This Study |  |  |
|  | Jacksonville, FL | 1 | t002 | SAMN15373856 | orfX-JI -ACME-SCCmecIIa |  |
|  | Wayne State University, Detroit, MI | 1 |  | #<br>SAMN35839406 |  |  |
| **2019** | Tampa General Hospital, Tampa, FL | 13 | t002(n=4), t062(n=8), t010(n=1) | This Study |  |  |
|  | University of Florida, Gainesville, FL | 2 | t002, t002 | PRJNA839096 | orfX-JI -ACME-SCCmecIIa |  |
| **2021** | Wayne State University, Detroit, MI | 1 | t045 | SAMN38637704 | SCCmecIIa |  |
| **2022** | Tampa General Hospital, Tampa, FL | 12 | t002(n=3), t062(n=7), t010(n=2) | This Study |  |  |
| **2024** | Tampa General Hospital, Tampa, FL | 1 | t062 | Unpublished |  |  |

### TGH-ST3390 Populations Have Three Different *spa*-types

From the 36 TGH-ST3390 strains we identified three different *spa*-types: t062 (n=18), t002 (n=11) and t010 (n=7). The ancestral *spa*-type t002 is by far the most common within the wider CC5 to which ST3390 belongs [29], however it is only the second most common type amongst TGH-ST3390 strains. While t010 and t062 are still associated with CC5, these *spa*-types are less frequently found according to the Ridom SpaServer. When reviewing the genomes of ST3390 isolates outside TGH we find that within the USA, absent the 2015-2016 strains from Chicago, all have t002, with the exception of a t045 isolate each from Baltimore and Detroit. Interestingly, when analyzing the Chicago strains, we find t002, t045 and t062 present, alongside 9 different *spa*-types that are unique only to this cluster. Despite this, these strains show a high level of similarity and cluster together in a distant clade from the rest of ST3390. Finally, the two international strains both have rare *spa*-types: the Canadian strain being t579 and the Dutch strain being t2065.

### ST3390 Strains Do Not Possess Typical SCCmec Clusters

When exploring genetic relatedness within TGH-ST3390 strains **(Figure 2A)**, we note all are *mecA*+, however the SCCmec cluster that they carry is highly unusual (**Figure 2B, Supplemental Table S1**). All possess SCCmecIIa, but, absent 4 individual isolates, all TGH-ST3390 strains also have additional genetic information in this region. For example, 24 strains also have a 6.2kb ACME II *arc* cluster with 99.9% homology to the *arc* cluster from USA300 ACME I, followed by a truncated 6.1kb joining region I (JI) from SCCmecIa. Furthermore, 3 isolates have the *arc* cluster–SCCmecIIa composite without the JI region. Additionally, there are 5 isolates that have an 11.5kb JI region from SCCmecVIII inserted at the *att* site of *orfX* instead of the ACME-JI composite. Outside of TGH we identified the ACME-JI-SCCmecIIa composite in 15/45 strains. A single isolate from Massachusetts appears to have the ACME-SCCmecIIa composite lacking the truncated SCCmecIa JI region. The SCCmecVIII JI-SCCmecIIa composite was not identified in any of the non-TGH strains, indicating this conformation is specific to the strains circulating within TGH.

**Figure 2.**
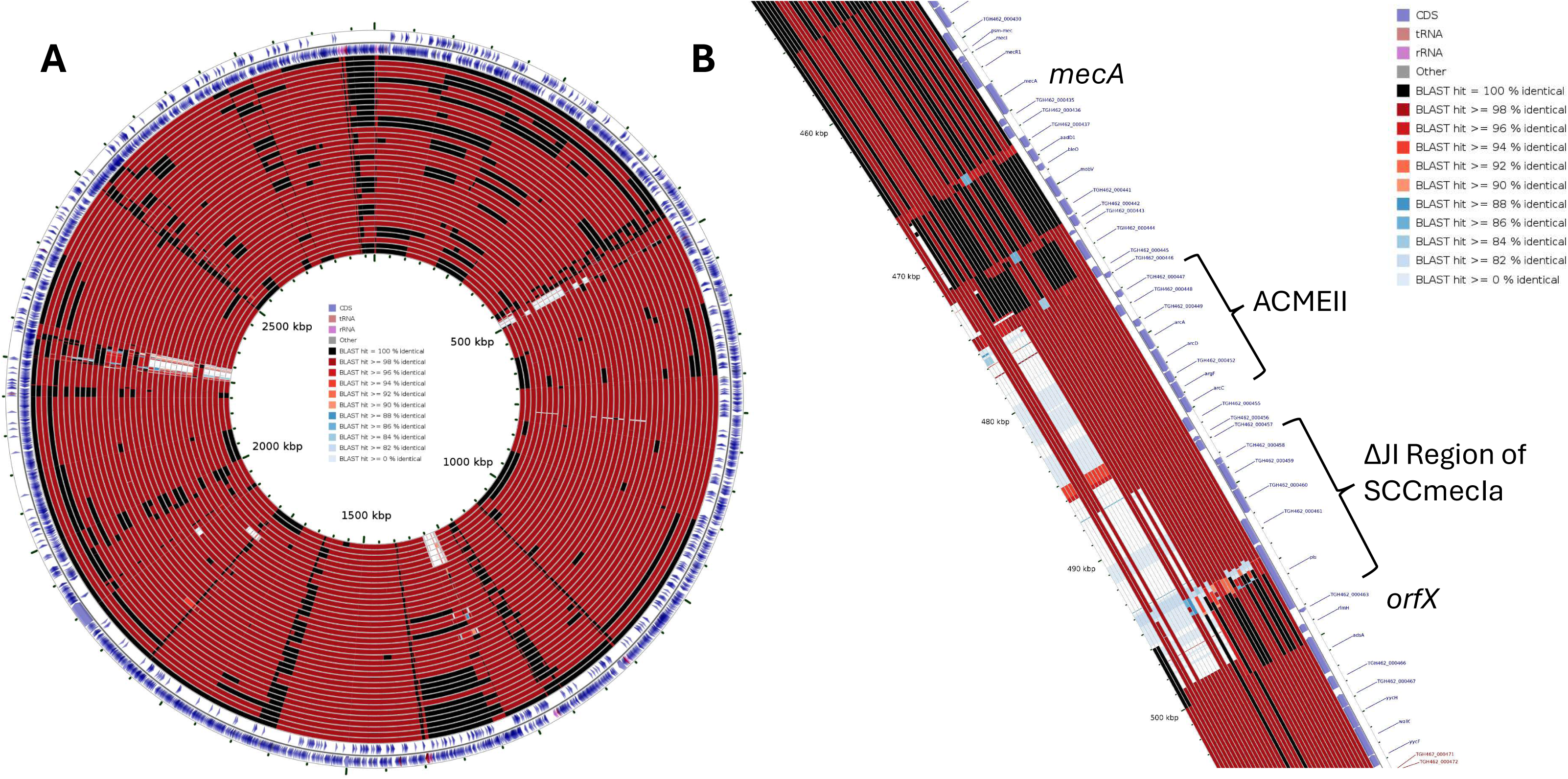
Homology Analysis of TGH-ST3390 Strains. **(A):** Blast maps of all 36 TGH-ST3390 using strain TGH462 as the root. Low identity genome regions are shown in blue or white. **(B):** An enlargement of A to show structural variation within SCCmec.

### Variations in the TGH ST3390 Mobilome

Using our genomic relatedness map we note that TGH-ST3390 strains have 4 prophage regions (**Figure 3A**). The first is a phiSa1int family member that is conserved within all TGH-ST3390 strains. The second is from the phiSa2int family, with its closest match being phi2958PVL. This region is highly conserved being present in 35/36 of TGH strains. Importantly, the phage is missing *lukF-PV*/*lukS-PV*, which has been observed in several other non-USA300 strains[12, 30]. Next is a member of the phiSa3int family, inserted into the beta-hemolysin gene. Interestingly this phage was present in 30 TGH-ST3390 strains but excised in 6; which is common during infection and antibiotic treatment and results in restored beta hemolytic activity. The final region is the least conserved and is missing in 13 strains. It’s a member of the phiSa7int family and its best match is to φJB. A study looking at the transduction efficiency found that φJB was an order of magnitude better at transducing resistance plasmids than other efficiently transducing phages[31].

**Figure 3.**
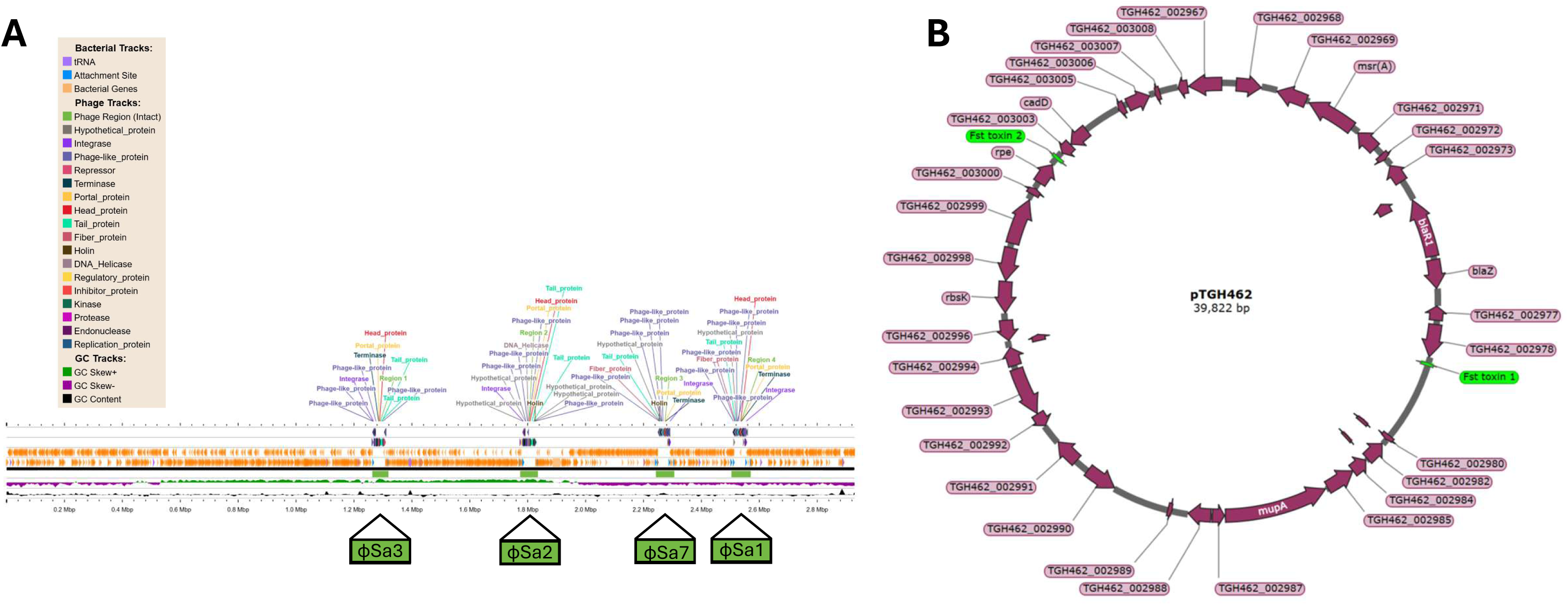
Analysis of Variable Genomic Elements within TGH-ST3390 Strains. **(A):** PHASTEST phage region map. Annotated in green are the different phage regions found in TGH462. **(B):** Plasmid map of pTGH462, highlighted in green are the two fst toxin genes.

### Mupirocin Resistance is Seemingly Maintained in TGH-ST3390 Strains via Plasmid Addiction

Within the population of TGH-ST3390 isolates, 24/36 strains carry the *mupA* gene on a plasmid, which drives high-level mupirocin resistance [32]. Importantly, carriage of *mupA* within the population continues without mupirocin selective pressure as TGH discontinued use of this antibiotic for decolonization in 2020. Using long read sequencing data, we detect a 39,822bp *mupA* plasmid in strain TGH462 (**Figure 3B**) that encodes two different Type 1 Toxin/Antitoxin (TA) fst family toxins. Importantly, in the context of plasmids, Type 1 TA systems are associated with ensuring inheritance via plasmid addiction [33–35]. In pTGH462, the two toxins contain a different P/D/S/TXXXG(C) motif within their transmembrane domain[36]. Fst toxin 1 is 30 aa long and encoded 4.7kb downstream from *mupA*; bearing the motif most common to this family, PXXXG. Fst toxin 2 is 33 aa long and is encoded 14.7kb upstream of *mupA.* It has a less common TXXXG motif, which is associated only with Staphylococcal species. Fst toxin 1 sequence is not unique to mupirocin plasmids, however, as it is also present in 6 *mupA*-TGH-ST3390 genomes, as well as all 24 of the *mupA*+ strains. Conversely, Fst toxin 2 was found in 23/24 TGH-ST3390 *mupA* plasmids, and only in a single *mupA*-genome. Interestingly, this latter strain appears to have only the first half of pTGH462 located upstream of *mupA*. As such, this dual plasmid addiction system within *mupA* bearing TGH-ST3390 strains may be the reason that this mupirocin resistance is maintained within the population.

### ST3390 Strains are Apigmented

When growing TGH-ST3390 strains we observed that all isolates were white in color, lacking the golden color that gives *S. aureus* its species name (**Figure 4A**). Pigment production in *S. aureus* is driven by the carotenoid biosynthesis operon, *crtOPQMN*. When reviewing genomes, we note that all of TGH-ST3390 strains have a 6aa in-frame deletion within the FAD/NAD(P)-binding domain of staphyloxanthin biosynthesis protein dehydrosqualene synthase (CrtN) (**Figure 4B**). To explore if this deletion is causative for apigmentation we amplified *crtOPQMN* from TGH993, cloned it into a shuttle vector and introduced it into a USA300-JE2 *crtN* mutant. In so doing, we observed that the TGH-3390 *crt* operon was unable to rescue the apigmented phenotype of the *crtN* mutant (**Supplemental Figure S1**), indicating that the TGH-3390 *crtN* gene is non-functional. When exploring genomes of the non-TGH ST3390 isolates we note that the same 6aa deletion is present in all strains.

**Figure 4.**
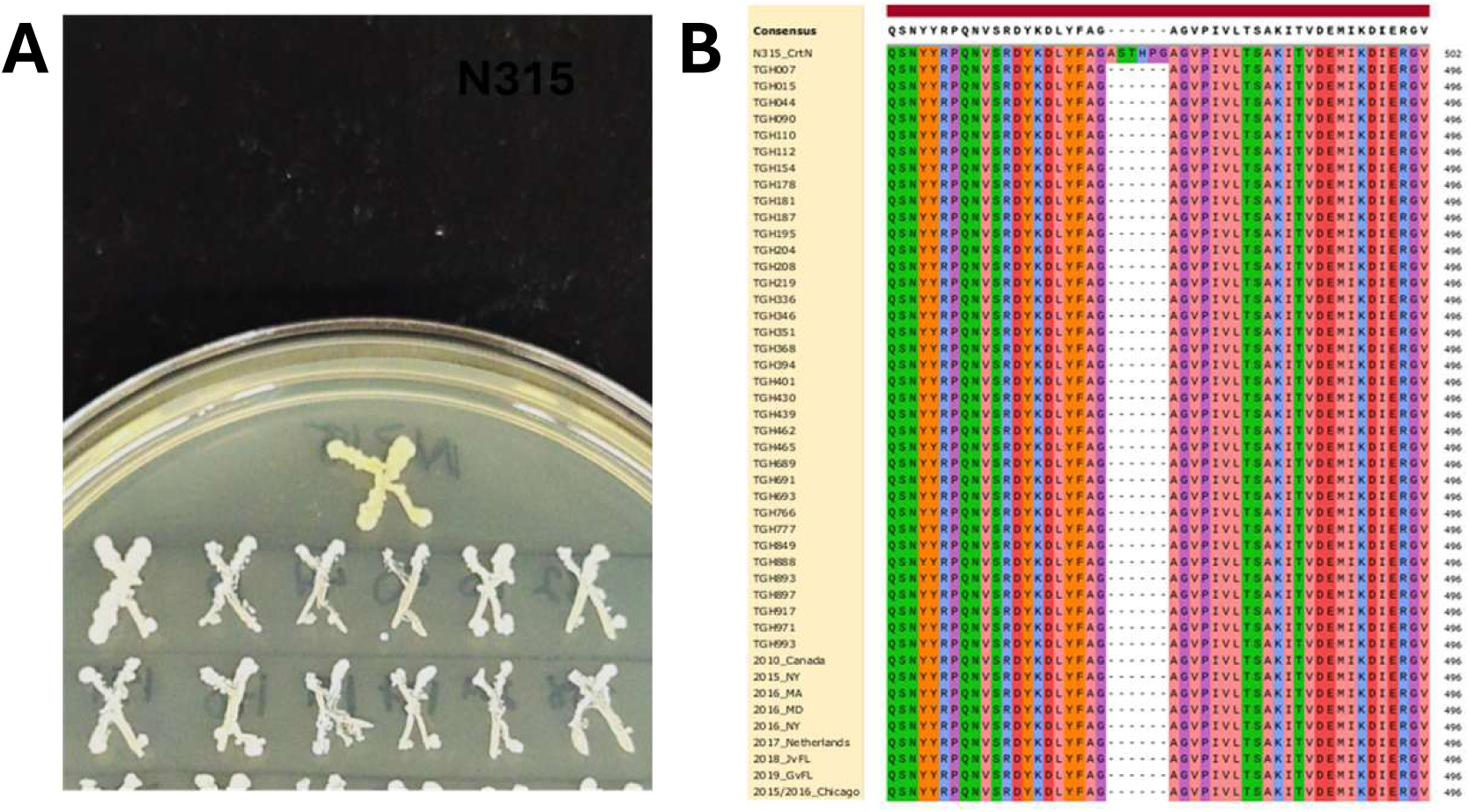
ST3390 Strains Lack the Characteristic Golden Staphyloxanthin Pigment of *S. aureus*. **(A):** TGH-ST3390 strains were patched on to Tryptic Soy Agar alongside the control CC5 strain N315 (labelled). (**B**): Protein alignment of CrtN from the CC5 control strain N315 compared to TGH-ST3390 strains. Also included are sequences from a single isolate for each year/city outside of TGH.

### Increased Blood Survival is Inversely Correlated with the Secreted Lytic Activity of TGH-ST3390 Strains

To phenotypically explore virulence capacities of TGH-ST3390 strains, we began by assessing their survivability in whole human blood (**Figure 5A**). These assays were performed alongside the well characterized ST5 isolate N315 to provide a baseline for the wider CC5 group. In so doing, we noted that the TGH-ST3390 population displayed a clear divide, with 11 isolates mirroring N315, with limited viability in blood, whilst the remaining 25 displayed robust survivability. When we expanded these studies to measure three primary virulence factor activities – hemolysis, proteolysis and cytolysis – we found an inverse correlation with blood survival (**Figure 5B, Supplemental Figure S2**). Here, most strains with limited viability in human blood were more likely to be highly proteolytic, hemolytic or cytolytic. Indeed, none of the strains that had robust survival in blood were highly proteolytic, and only one was highly hemolytic and cytolytic (TGH394).

**Figure 5.**
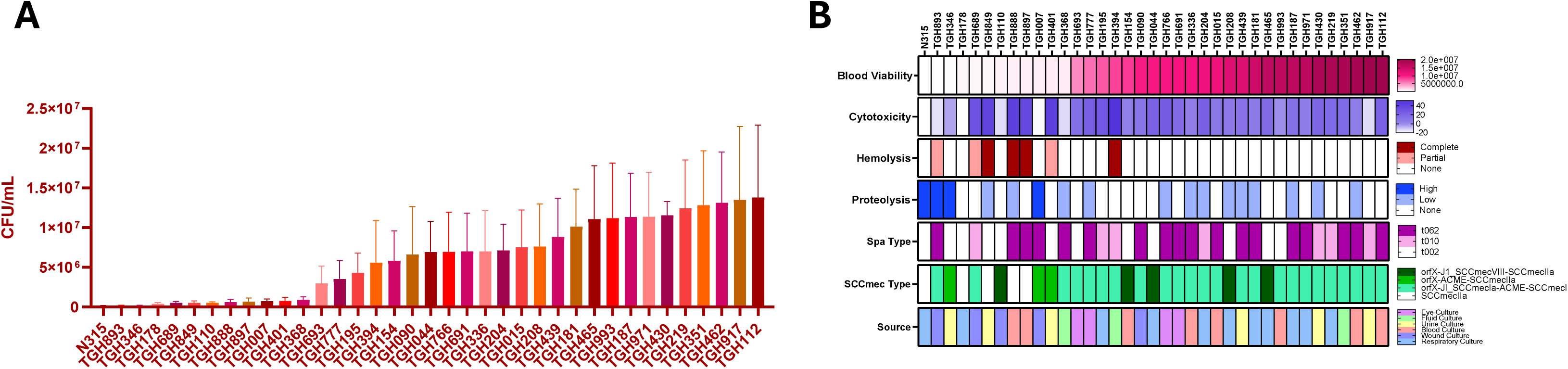
Secreted Lytic Activity of TGH-ST3390 Strains is Inversely Correlated with Survival in Human Blood. **(A):** Strains were grown overnight in TSB, before being standardized to an OD_600_ of 0.05 and inoculated into whole human blood. Strains were incubated for 6h before viability was determined via plating. Error bars are ±SEM; n=3. **(B):** Composite representation of secreted lytic activities for all strains, alongside spa-type, SCCmec Type and source of origin. More details on lytic activity assays can be found in Supplemental Figure S2.

### TGH-3390 strains Have Enhanced Virulence in a Murine Model of Sepsis

To determine the virulence of TGH-ST3390 strains we next used a murine model of sepsis. Here we chose a representative strain from each of the three *spa*-types, alongside the comparator CC5 strain N315, and used them to infect mice. When assessing the mortality of animals during the infection period, no mice infected with N315 or TGH178 (t002) died (**Figure 6A**). In contrast, two mice infected with TGH112 (t062) and three mice infected with TGH219 (t010) died. Those mice surviving the infection period were euthanized and the kidneys and liver were harvested. When these were plated for viability (**Figure 6BC**) we observed similar trends to our mortality data. Specifically, TGH112 had 9.91-fold greater burden in the kidneys and 9.87-fold greater burden in the liver, compared to N315. For TGH219, it had 16.6-fold greater burden in the kidneys and 516.6-fold greater burden in the liver, compared to N315. Perhaps more strikingly, the kidneys from TGH112 and TGH219 infected mice had greater signs of disease severity, including alterations in color and size of the organs (compared to N315), and a greater accumulation of abscesses.

**Figure 6.**
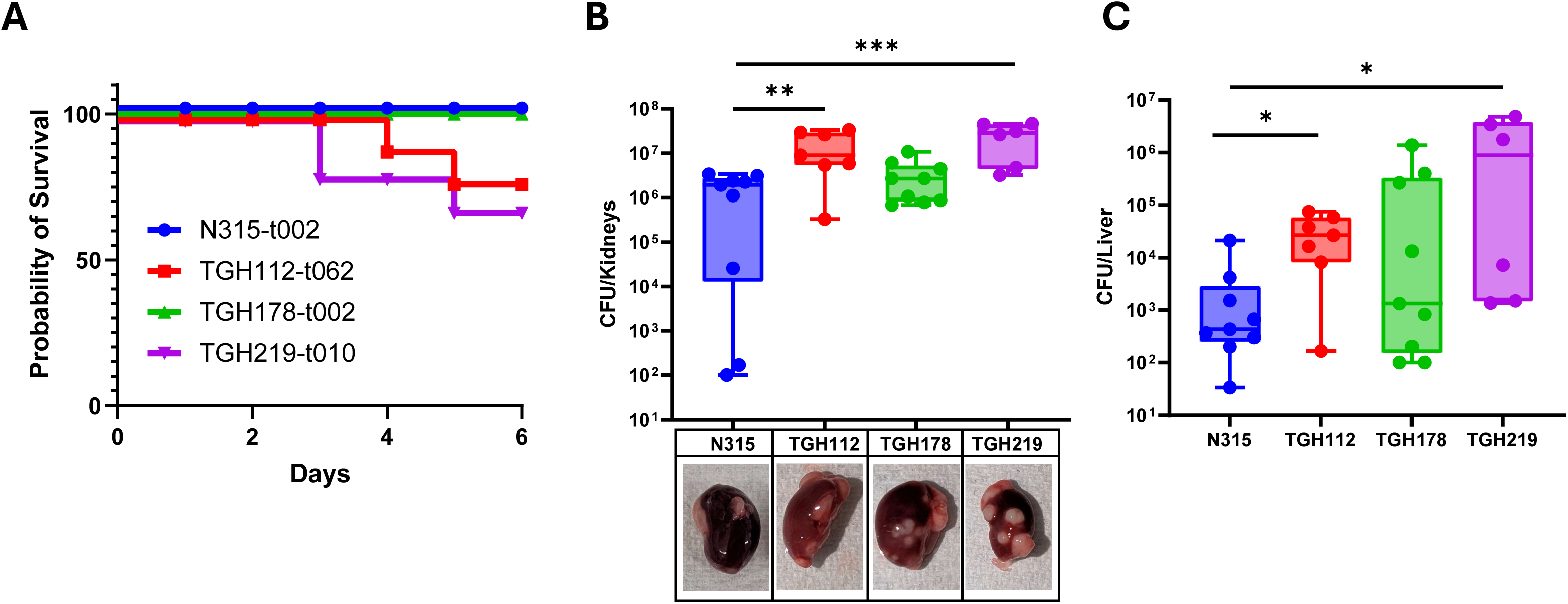
TGH-ST3390 Strains With *spa*-Types t010 and t062 Have Enhanced Virulence in a Murine Model of Sepsis. **(A):** Nine female CD-1 mice were inoculated via tail vein with 1x10^8^ of the strains shown. Infections were allowed to proceed for 7 days and the mortality of mice was recorded. For mice that survived the infection period, the **(B)** kidneys or **(C)** liver were harvested, homogenized, and plated for viability. For (B) inset are representative images of kidneys from surviving mice. Statistical significance was determined using a Log Rank (A) or Mann Whitney Test (B-C). * = p< 0.05; ** = p< 0.01, *** = p< 0.001.

## Discussion

Herein we present the identification and characterization of a rare *S. aureus* Sequence Type, ST3390. The appearance of STT3390, first in Canada, and then along the Eastern, mirrors that seen with other CC5-ST5 descendant clones. For example, in Brazil, clone ST105 evolved from ST5, has a higher propensity towards blood stream infections, and is better at evading phagocytosis and killing by monocytes than other CC5 isolates [13]. Similarly, in Japan and China, ST764 is an offshoot of ST5 and causes severe invasive disease with enhanced virulence over ST5 strains [12, 37]. We thus suggest that ST3390 can be seen as a North American expansion of CC5 with newly develop virulence properties. Interestingly, ST3390 mirrors ST764 at least in part as it has acquired components of ACME via horizontal gene exchange.

Amongst ST3390 isolates there is a relatively good *spa*-typing consensus, at least within the TGH population, of t062 > t002 > t010. Outside of a single isolate in Detroit and Baltimore, and the Chicago outbreak, USA strains only possess t002, although the two non-US isolates also diverge. The Chicago strain set is fascinating as it contains 12 different *spa*-types across 34 strains. This kind of variation within a ST is not uncommon, as, for example, ST5s have multiple *spa*-types [38]. One of the reasons that *spa* sequences are so variable is because they are formed by patterns of tandem sequence repeats that are subject to frequent polymerase staling during DNA replication and therefore evolve more quickly than most protein-coding regions of the genome [39]. It also is possible that variant *spa*-types are selected by the host immune system during infection because Spa is a secreted virulence factor [40–42]. Thus, it may be that t002 is the standard within ST3390, but that this is subject to geographical variation/specialization during outbreaks and expansion events. Indeed, our most virulent strain is type t010, which is only found at TGH.

We also report several different composite SCCmec elements in ST3390 strains. These arise either via recombination events between different SCCmec types or through the addition or deletion of mobile genetic elements [6, 43]. Composite elements also potentially arise more readily in hospital-associated strains due to horizontal gene transfer between diverse species as well as the selective pressures from antibiotics[44]. We note that the majority of TGH-ST3390 isolates harbor a truncated form of ACME II and modified JI region from SCCmecIa, located between *orfX* and SCCmec. Whilst the ACMEII *arc* cluster bears high similarity to that of ACME I from USA300 strains, ACME II is primarily identified with *S. epidermidis* ATCC 12228. Furthermore, the *ccr* genes from SCCmecVIII (and found in some TGH-ST3390 strains) are nearly identical to those found in *S. epidermidis*. This indicates the potential for ST3390 to readily attain genetic features not only from other *S. aureus* strains but also from other Staphylococcal species. It is potentially noteworthy that others have shown that the unique profiles of combinatorial SCCmec elements have the potential to confer enhanced virulence and antimicrobial resistance [45, 46], both key features of TGH-ST3390 strains.

The lack of pigmentation in ST3390 strains is of significant interest. It has previously been reported that apigmented *S. aureus* strains have impaired neutrophil survival, are more sensitive to reactive oxygen species, and have limited virulence in murine models [47]. Strikingly, we see quite the opposite with TGH-ST3390 strains, which display increased neutrophil killing and enhanced virulence during sepsis compared to the pigment CC5 isolate N315. Interestingly, recent studies using naturally apigmented lineages of *S. aureus* (including ST20 and ST25) demonstrate that there is actually no difference in the virulence of these strains compared to pigmented counterparts [38], which mirrors our findings. Another interesting consideration is that a majority of naturally occurring apigmented *S. aureus* strains lack the *crtOPQMN* operon [38, 48], whereas ST3390 merely has a conserved deletion with *crtN*. This perhaps suggests the relatively recent acquisition of this mutation and consequent apigmented phenotype. It will be of significant interest to determine if ST3390 strains can have their pigmentation restored via repair of the *crtN* gene, and how this influences strain behavior.

When considering our laboratory-based assessment of virulence, our findings at first glance may appear contradictory – in that we find those strains that are highly proteolytic, cytolytic and hemolytic have limited survivability in human blood. This does, however, make sense, as we know that strains of *S. aureus* that are hyperproteolytic are typically attenuated in infection models [49]. This is driven by a role for these enzymes in controlling the stability of self-derived surface associated and secreted virulence factors [50]. Thus, the overproduction of secreted proteases limits the abundance of other virulence factors required for survival in, for example, human blood [51]. Similarly, we know that limiting the hemolytic or cytolytic capacity of *S. aureus* can have a positive impact on bacterial survival during infection [52, 53]. Thus, it appears that virulence factor expression in ST3390 strains appears to have achieved an ideal equilibrium to facilitate survival and dissemination within the host.

In conclusion, we report the characterization of a rare CC5 descendant clone with the capacity for hypervirulence that has taken root in the Tampa Bay region. We first detected it during collection for this project in 2017 and have found new strains each year we sample. We believe the number of ST3390 strains from our hospital is far greater than that reported herein – we have found many white *S. aureus* isolates in our collection that display high level mupirocin resistance, indicating that they are quite likely to be ST3390 strains. Given that the Tampa Bay metro area has >3 million residents and that ST3390 strains are found in multiple cities in Florida and the wider USA, there is a clear and pressing need to further characterize this novel lineage. A key consideration is whether this will ultimately resolve in time, or if we are perhaps witnessing an expansion (locally or beyond) of this ST akin to that seen for ST8-USA300 in the 1990s.

## Acknowledgements

This study was supported by grants AI124458 and AI157506 (both to L.N.S.) from the National Institute of Allergy and Infectious Diseases and the USF Provost’s CREATE award (LNS and KK).

## References

1. David MZ, Daum RS. Community-Associated Methicillin-Resistant *Staphylococcus aureus*: Epidemiology and Clinical Consequences of an Emerging Epidemic. Clinical Microbiology Reviews 2010; 23:616–87.

2. Chambers HF, Deleo FR. Waves of resistance: Staphylococcus aureus in the antibiotic era. Nat Rev Microbiol 2009; 7:629–41.

3. Harmsen D, Claus H, Witte W, et al. Typing of Methicillin-Resistant *Staphylococcus aureus* in a University Hospital Setting by Using Novel Software for *spa* Repeat Determination and Database Management. Journal of Clinical Microbiology 2003; 41:5442–8.

4. Classification of Staphylococcal Cassette Chromosome *mec* (SCC *mec* ): Guidelines for Reporting Novel SCC *mec* Elements. Antimicrobial Agents and Chemotherapy 2009; 53:4961-7.

5. Lakhundi S, Zhang K. Methicillin-Resistant Staphylococcus aureus: Molecular Characterization, Evolution, and Epidemiology. Clinical Microbiology Reviews 2018; 31.

6. Hanssen A-M, Ericson Sollid JU. SCC *mec* in staphylococci: genes on the move. FEMS Immunology & Medical Microbiology 2006; 46:8–20.

7. Thurlow LR, Joshi GS, Richardson AR. Virulence strategies of the dominant USA300 lineage of community-associated methicillin-resistant *Staphylococcus aureus* (CA-MRSA). FEMS Immunology & Medical Microbiology 2012; 65:5–22.

8. Dyzenhaus S, Sullivan MJ, Alburquerque B, et al. MRSA lineage USA300 isolated from bloodstream infections exhibit altered virulence regulation. Cell Host & Microbe 2023; 31:228–42.e8.

9. Thiede SN, Snitkin ES, Trick W, et al. Genomic Epidemiology Suggests Community Origins of Healthcare-Associated USA300 Methicillin-Resistant *Staphylococcus aureus*. The Journal of Infectious Diseases 2022.

10. Sola C, Paganini H, Egea AL, et al. Spread of Epidemic MRSA-ST5-IV Clone Encoding PVL as a Major Cause of Community Onset Staphylococcal Infections in Argentinean Children. PLoS ONE 2012; 7:e30487.

11. Jian Y, Zhao L, Zhao N, et al. Increasing prevalence of hypervirulent ST5 methicillin susceptible *Staphylococcus aureus* subtype poses a serious clinical threat. Emerging Microbes & Infections 2021; 10:109–22.

12. Takano T, Hung W-C, Shibuya M, et al. A New Local Variant (ST764) of the Globally Disseminated ST5 Lineage of Hospital-Associated Methicillin-Resistant Staphylococcus aureus (MRSA) Carrying the Virulence Determinants of Community-Associated MRSA. Antimicrobial Agents and Chemotherapy 2013; 57:1589–95.

13. Viana AS, Nunes Botelho AM, Moustafa AM, et al. Multidrug-Resistant Methicillin-Resistant *Staphylococcus aureus* Associated with Bacteremia and Monocyte Evasion, Rio de Janeiro, Brazil. Emerging Infectious Diseases 2021; 27:2825–35.

14. Viana AS, Tótola LPDV, Figueiredo AMS. ST105 Lineage of MRSA: An Emerging Implication for Bloodstream Infection in the American and European Continents. Antibiotics 2024; 13:893.

15. Virgillio AN and Felton EA JJ, Kennedy S, Becker DN, Lima A, Atrubin K, Cella E, Azarian T, Silbert S, Shaw LN, Kim, K. Genomic and Phenotypic Characterization of Mupirocin Resistant Staphylococcus aureus Clinical Isolates. 2025.

16. Biguenet A, Bordy A, Atchon A, Hocquet D, Valot B. Introduction and benchmarking of pyMLST: open-source software for assessing bacterial clonality using core genome MLST. Microb Genom 2023; 9.

17. Petit RA, 3rd, Read TD. Bactopia: a Flexible Pipeline for Complete Analysis of Bacterial Genomes. mSystems 2020; 5.

18. Wishart DS, Han S, Saha S, et al. PHASTEST: faster than PHASTER, better than PHAST. Nucleic Acids Res 2023; 51:W443–W50.

19. Stothard P, Wishart DS. Circular genome visualization and exploration using CGView. Bioinformatics 2005; 21:537–9.

20. Bose JL, Fey PD, Bayles KW. Genetic Tools To Enhance the Study of Gene Function and Regulation in Staphylococcus aureus. Applied and Environmental Microbiology 2013; 79:2218–24.

21. Gimza BD, Marroquin SM, Shaw LN. An Ex Vivo Model for Assessing Growth and Survivability of Staphylococcus aureus in Whole Human Blood. Methods Mol Biol 2021; 2341:127–31.

22. Dumont AL, Nygaard TK, Watkins RL, et al. Characterization of a new cytotoxin that contributes to Staphylococcus aureus pathogenesis. Molecular Microbiology 2011; 79:814–25.

23. Jobson M-E, Tomlinson BR, Mustor EM, et al. SSR42 is a Novel Regulator of Cytolytic Activity in *Staphylococcus aureus*: Cold Spring Harbor Laboratory, 2024.

24. Guthrie JL, Teatero S, Hirai S, et al. Genomic Epidemiology of Invasive Methicillin-Resistant *Staphylococcus aureus* Infections Among Hospitalized Individuals in Ontario, Canada. The Journal of Infectious Diseases 2020; 222:2071–81.

25. Tenover FC, Tickler IA, Le VM, Dewell S, Mendes RE, Goering RV. Updating Molecular Diagnostics for Detecting Methicillin-Susceptible and Methicillin-Resistant Staphylococcus aureus Isolates in Blood Culture Bottles. Journal of Clinical Microbiology 2019; 57.

26. Bispo PJM, Ung L, Chodosh J, Gilmore MS. Hospital-Associated Multidrug-Resistant MRSA Lineages Are Trophic to the Ocular Surface and Cause Severe Microbial Keratitis. Frontiers in Public Health 2020; 8.

27. Popovich KJ, Green SJ, Okamoto K, et al. MRSA Transmission in Intensive Care Units: Genomic Analysis of Patients, Their Environments, and Healthcare Workers. Clinical Infectious Diseases 2021; 72:1879–87.

28. Schouls LM, Witteveen S, Van Santen-Verheuvel M, et al. Molecular characterization of MRSA collected during national surveillance between 2008 and 2019 in the Netherlands. Communications Medicine 2023; 3.

29. Asadollahi P, Farahani NN, Mirzaii M, et al. Distribution of the Most Prevalent Spa Types among Clinical Isolates of Methicillin-Resistant and -Susceptible Staphylococcus aureus around the World: A Review. Frontiers in Microbiology 2018; 9.

30. Goerke C, Pantucek R, Holtfreter S, et al. Diversity of Prophages in Dominant *Staphylococcus aureus* Clonal Lineages. Journal of Bacteriology 2009; 191:3462–8.

31. Varga M, Pantu Ček R, Ru Žičková V, Doškař J. Molecular characterization of a new efficiently transducing bacteriophage identified in meticillin-resistant Staphylococcus aureus. J Gen Virol 2016; 97:258–68.

32. Patel JB, Gorwitz RJ, Jernigan JA. Mupirocin resistance. Clin Infect Dis 2009; 49:935–41.

33. Weaver KE, Reddy SG, Brinkman CL, Patel S, Bayles KW, Endres JL. Identification and characterization of a family of toxin-antitoxin systems related to the Enterococcus faecalis plasmid pAD1 par addiction module. Microbiology (Reading) 2009; 155:2930–40.

34. Kroll J, Klinter S, Schneider C, Voß I, Steinbüchel A. Plasmid addiction systems: perspectives and applications in biotechnology. Microbial Biotechnology 2010; 3:634–57.

35. Kwong SM, Jensen SO, Firth N. Prevalence of Fst-like toxin-antitoxin systems. Microbiology (Reading) 2010; 156:975–7.

36. Weaver K. The Fst/Ldr Family of Type I TA System Toxins: Potential Roles in Stress Response, Metabolism and Pathogenesis. Toxins 2020; 12:474.

37. Xiao Y, Han W, Wang B, et al. Phylogenetic analysis and virulence characteristics of methicillin-resistant *Staphylococcus aureus* ST764-SCC *mec* II: an emerging hypervirulent clone ST764-t1084 in China. Emerging Microbes & Infections 2023; 12.

38. Zhang J, Suo Y, Zhang D, Jin F, Zhao H, Shi C. Genetic and Virulent Difference Between Pigmented and Non-pigmented Staphylococcus aureus. Front Microbiol 2018; 9:598.

39. Nübel U, Roumagnac P, Feldkamp M, et al. Frequent emergence and limited geographic dispersal of methicillin-resistant *Staphylococcus aureus*. Proceedings of the National Academy of Sciences 2008; 105:14130–5.

40. Howden BP, Giulieri SG, Wong Fok Lung T, et al. Staphylococcus aureus host interactions and adaptation. Nature Reviews Microbiology 2023; 21:380–95.

41. Thammavongsa V, Kim HK, Missiakas D, Schneewind O. Staphylococcal manipulation of host immune responses. Nature Reviews Microbiology 2015; 13:529–43.

42. Falugi F, Kim HK, Missiakas DM, Schneewind O. Role of Protein A in the Evasion of Host Adaptive Immune Responses by Staphylococcus aureus. mBio 2013; 4:e00575–13-e.

43. Berglund C, Mölling P, Sjöberg L, Söderquist B. Predominance of staphylococcal cassette chromosome mec (SCCmec) type IV among methicillin-resistant Staphylococcus aureus (MRSA) in a Swedish county and presence of unknown SCCmec types with Panton-Valentine leukocidin genes. Clin Microbiol Infect 2005; 11:447–56.

44. Deurenberg RH, Vink C, Kalenic S, Friedrich AW, Bruggeman CA, Stobberingh EE. The molecular evolution of methicillin-resistant Staphylococcus aureus. Clinical Microbiology and Infection 2007; 13:222–35.

45. Urushibara N, Kawaguchiya M, Kobayashi N. Two novel arginine catabolic mobile elements and staphylococcal chromosome cassette mec composite islands in community-acquired methicillin-resistant Staphylococcus aureus genotypes ST5-MRSA-V and ST5-MRSA-II. Journal of Antimicrobial Chemotherapy 2012; 67:1828–34.

46. Sabat AJ, Köck R, Akkerboom V, et al. Novel Organization of the Arginine Catabolic Mobile Element and Staphylococcal Cassette Chromosome *mec* Composite Island and Its Horizontal Transfer between Distinct Staphylococcus aureus Genotypes. Antimicrobial Agents and Chemotherapy 2013; 57:5774–7.

47. Liu GY, Essex A, Buchanan JT, et al. Staphylococcus aureus golden pigment impairs neutrophil killing and promotes virulence through its antioxidant activity. J Exp Med 2005; 202:209–15.

48. Holt DC, Holden MT, Tong SY, et al. A very early-branching Staphylococcus aureus lineage lacking the carotenoid pigment staphyloxanthin. Genome Biol Evol 2011; 3:881–95.

49. Zielinska AK, Beenken KE, Mrak LN, et al. sarA-mediated repression of protease production plays a key role in the pathogenesis of Staphylococcus aureus USA300 isolates. Mol Microbiol 2012; 86:1183–96.

50. Gimza BD, Jackson JK, Frey AM, et al. Unraveling the Impact of Secreted Proteases on Hypervirulence in Staphylococcus aureus. mBio 2021; 12.

51. Kolar SL, Ibarra JA, Rivera FE, et al. Extracellular proteases are key mediators of Staphylococcus aureus virulence via the global modulation of virulence-determinant stability. Microbiologyopen 2013; 2:18–34.

52. Das S, Lindemann C, Young BC, et al. Natural mutations in a Staphylococcus aureus virulence regulator attenuate cytotoxicity but permit bacteremia and abscess formation. Proc Natl Acad Sci U S A 2016; 113:E3101–10.

53. Laabei M, Uhlemann A-C, Lowy FD, et al. Evolutionary Trade-Offs Underlie the Multi-faceted Virulence of Staphylococcus aureus. PLOS Biology 2015; 13:e1002229.

