## Supplemental Figures 1-2 for "Emergence of ST3390: A non-pigmented HA-MRSA clone with enhanced virulence"

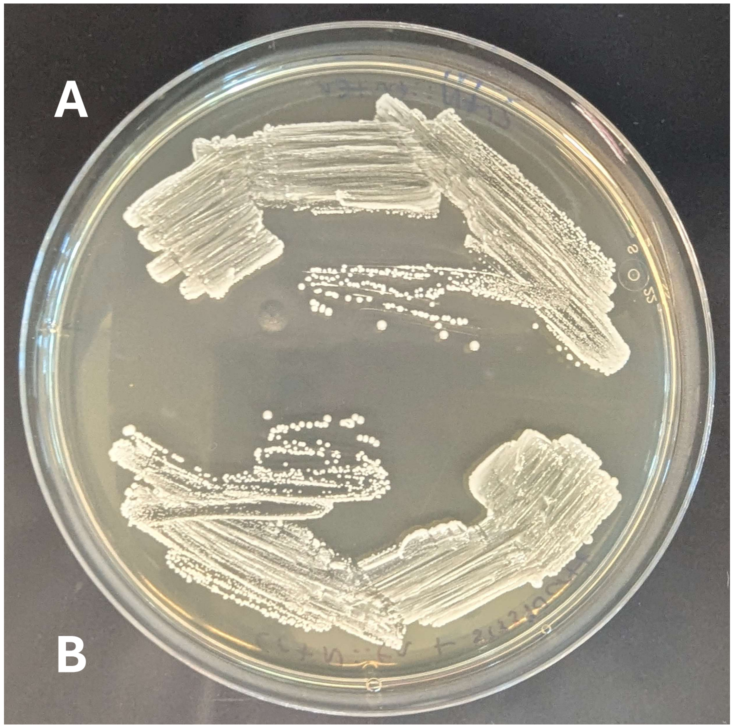


**Supplemental Figure S1. The CrtN Protein of ST3390 Strains is Defective.** A USA300 Je2 crtN transposon mutant bearing an empty pMK4 shuttle vector was streaked onto Tryptic Soy Agar (**A**). Shown in **B** is the same strain, however the pMK4 vector contained the entire *crtOPQMN* operon from TGH-ST3390 strain TGH993 under control of its native promoter.

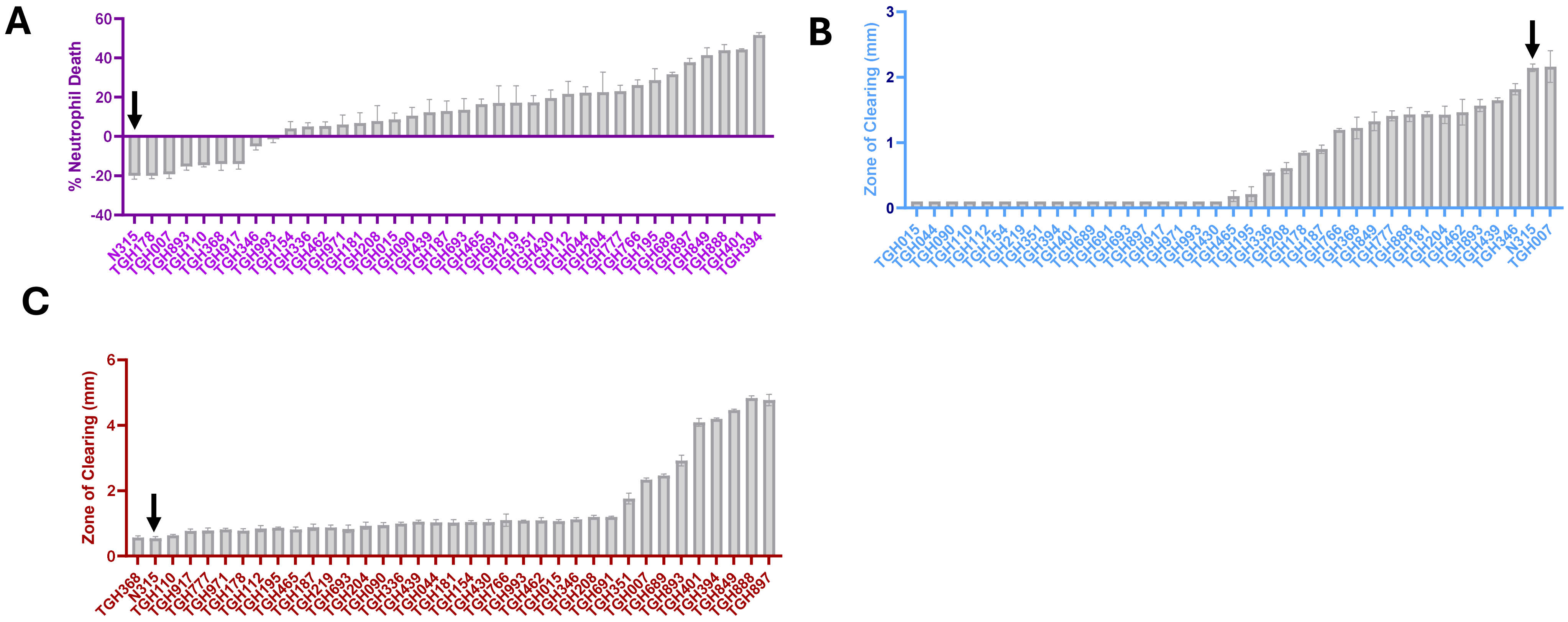


**Supplemental Figure S2. Secreted Lytic Activities for TGH-ST3390 Strains. (A):** Cytolysis of neutrophils designated as % death. **(B):** Proteolytic activity measured on 5% casein agar. **(C):** Hemolysis on 5% sheep blood. Error bars are ±SEM; n=3. The comparator CC5-ST5 strain N315 is highlighted in each panel via a black arrow.
